# Differential tissue stiffness of body column facilitates locomotion of *Hydra* on solid substrates

**DOI:** 10.1101/2020.02.29.971044

**Authors:** Suyash Naik, Manu Unni, Devanshu Sinha, Shatruhan Singh Rajput, P. Chandramouli Reddy, Elena Kartvelishvily, Inna Solomonov, Irit Sagi, Apratim Chatterji, Shivprasad Patil, Sanjeev Galande

## Abstract

The bell-shaped members of Cnidaria typically move around by swimming, whereas the *Hydra* polyp can perform locomotion on solid substrates in aquatic environment. To address the biomechanics of locomotion on rigid substrates, we studied the ‘somersaulting’ locomotion in *Hydra*. We applied atomic force microscopy to measure the local mechanical properties of *Hydra’s* body column and identified the existence of differential Young’s modulus between the shoulder region versus rest of the body column at 3:1 ratio. We show that somersault primarily depends on differential tissue stiffness of the body column and is explained by computational models that accurately recapitulate the mechanics involved in this process. We demonstrate that perturbation of the observed stiffness variation in the body column by modulating the extracellular matrix (ECM) polymerization impairs the ‘somersault’ movement. These results provide mechanistic basis for the evolutionary significance of differential extracellular matrix properties and tissue stiffness.

## Introduction

Locomotion enables organisms to move from one place to another. The need to move around evolved very early in life forms, which accorded several advantages to the organisms. Locomotion offers exploration opportunities for food and water source, mates, niche and more importantly, escaping predators. Morphological diversities and tissue type innovations facilitated in the evolution of various means of locomotion in different organisms. The organisms that use specialized appendages and body oscillations to propel themselves are often known to share common biomechanical principles (Biewener, 1990, Gray, 1933, Alexander, 2003) While unicellular organisms use dedicated organelles such as cilia, flagella or pseudopodia, the multicellular organisms exhibit a complex system of coordination among specific cell types to achieve locomotion (Bray, 2000). Although organisms from phyla such as Ctenophora and sponges are capable of locomotion involving coordination of their multicellular body, they do not possess specialized tissues unlike the more complex eumetazoans (Bond and Harris, 1988, Matsumoto, 1991). Basal metazoans evolved two types of locomotion - fluid-dependent and substrate-dependent. Cnidarians have acquired the ability to perform coordinated locomotion both in water and solid substrates and are the earliest phylum to evolve differentiated neuronal/muscular tissues and extracellular matrix properties (Bode, 1996, Bode et al., 1990, Dupre and Yuste, 2017).

Cnidarians typically have two different types of body structures viz. the medusa and the polyp forms (Galliot, 2000). The medusal organisms such as Jellyfish perform the movement with the help of thrust or pull of fluids using the umbrella, and hence are dependent on the mechanics of fluids (Gemmell et al., 2015, Anderson and DeMont, 2000). The direction of movement can be controlled by directing the fluid flow with the help of muscles (Anderson and DeMont, 2000). *Hydra* is a cnidarian polyp lacking the medusal stage in its life cycle. It has a very slender yet extremely flexible body column having equally flexible tentacles at the oral end. The flexibility of *Hydra* is primarily due to its unique extracellular matrix composition (Deutzmann et al., 2000). In the case of *Hydra*, the more complex modes of locomotion compared to floating, utilize the neuro-muscular system. The muscular cells in the two germ layers are oriented orthogonally with respect to each other, which help in controlling both the length as well as the radial width of the organism (Aufschnaiter et al., 2017). These muscular cells are controlled by different neural circuits to achieve a range of coordinated behavioural activities (Dupre and Yuste, 2017, Han et al., 2018, Davis et al., 1968). This ability of *Hydra* to coordinate contraction and relaxation helps the polyp to perform a range of movements such as swaying, looping and somersaulting (Trembley, 1744).

In the current study, we have focused on *Hydra’s* somersault, which involves coordinated movements of different body parts. It includes adhering to the substrate with the tentacles for traction. The body is then moved around the head in a semi-circular arc with characteristic contractions and relaxations (Figure 1). It is initiated by stretching the body column almost to double its length (stage 1, Figure 1) and by attachment of the head to the substratum with the help of tentacles and hypostome (Trembley, 1744, Han et al., 2018). During this process, the region of body column just below the head referred to here as ‘shoulder’, is bent by almost 90° angle. The basal disc at the bottom of the foot is then released, relaxing the stretched body column and accentuating the bend in the shoulder (stages 2-3). The bend straightens out (or is released) to achieve ‘upside down’ position of the body column perpendicular to the substratum (stages 4-6) (Figure 1, Movie 1). The process is completed by bending the body column, followed by attachment of basal disc to a new position. Finally, the oral end pushes itself away with the help of tentacles, and the body attains the upright position.

**Figure 1:**
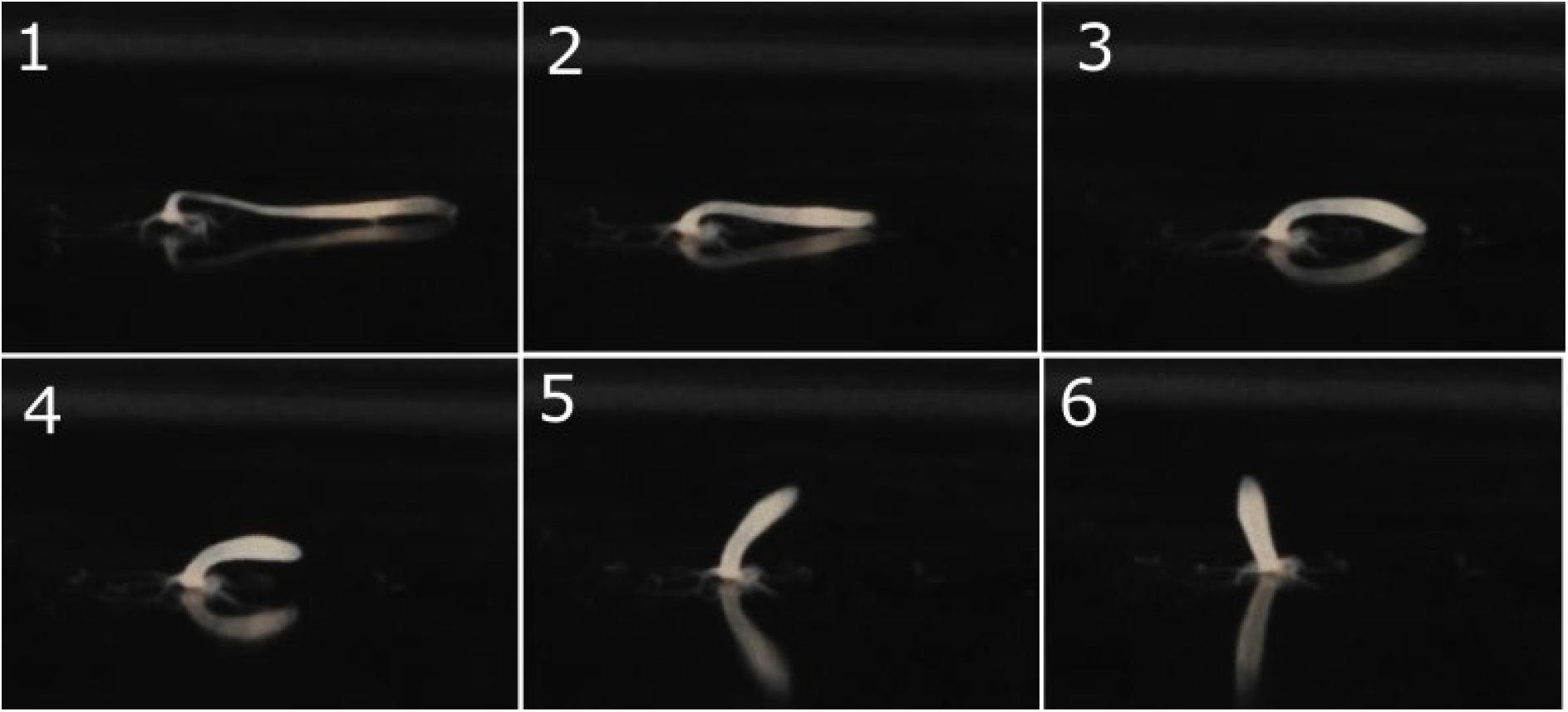
The *Hydra* Somersault. The stages in the part of somersault movement of *Hydra*. In stage 1, the body column is stretched, and the tentacles hold onto the substrate. In stage 2, the basal end is released. In stage 3, the body column contracts. In stage 4 and 5 the body column is lifted.

To understand biomechanics governing the somersault, we produced a spatially-resolved elasticity profile of *Hydra’s* body column using an Atomic Force Microscope (AFM). AFM allows micro-elasticity maps of biological materials (Tao et al., 1992). The profile obtained using AFM experiments is used to model *Hydra’s* body column in computer simulations to recapitulate part of the somersault movement. Further, we performed biological tests on *Hydra’s* ability to somersault if the differential tissue stiffness is removed by mechanical and chemical means. We observed that polyps with uniform stiffness along the body column lose their ability to perform a somersault.

Although the somersault is not the same as walking, there are interesting parallels in terms of the forces and energy recycling to reduce muscular work. The observed differential in tissue stiffness is possibly a harbinger in the evolution of a spatially heterogeneous stiffness of extracellular matrix which enables movement. The extracellular matrix-mediated stiffness differential has presumably been an ancient mechanism to accomplish specialized tissue function. In this study, we show that the extracellular matrix assumes a pseudo-skeletal property to help *Hydra* pull off the somersaulting stunt.

## Results

### Measurement of local variation in tissue stiffness

Multicellular organisms use spatially differentiated tissue types, such as exoskeleton in invertebrates or musculoskeletal systems as seen in vertebrates, to resist strain and sustain tractions during locomotion (Biewener, 1990, Dickinson et al., 2000). These specialized tissues exhibit differential stiffness properties, which help in locomotion. It was not known if *Hydra* possesses any tissue elasticity-dependent mechanisms to aid in its locomotion. We used Atomic Force Microscopy (AFM) for the first time to produce a spatially resolved map of tissue elasticity in *Hydra*. As shown in Figure 2A, we attached glass beads (diameter 25 μm) to cantilevers and carefully allowed the bead to touch the *Hydra* body laid on a BSA coated coverslip. With help of servo control, we record the force curves in which the load on the tissue and its deformations are measured by recording the cantilever deflections and substrate displacements. Such force curves are taken at locations separated by 100 μm along the body column. At each location we collect 25 force curves over a 5×5 grid and the area of 25 μm x 25 μm. A total of three polyps were used (N=3) in each experiment for stiffness measurement. Using Hertz contact mechanics, we estimated the Young’s modulus from each force curve, and the average was then calculated for each location (For further details see Figure S1 A and B). Figure 2B shows a typical force curve and fit using the Hertz model. Figure 2C shows the elasticity profile acquired using these measurements. We observed higher stiffness in the region below the base of tentacles, which extends up to 25 per cent of the length of the body column, referred to as the shoulder region. The shoulder region is nearly three-fold stiffer (Y =1480 ± 20 Pa) compared to the rest of the body (Y = 450 ± 6 Pa). For tentacles, the Young’s modulus (Y) is 378 ± 11 Pa (Figure S1C). These measurements revealed a steep drop in stiffness at the junction of the shoulder and the rest of the body column (Figure 2C, Figure S1D). The entire set of measurements was repeated over three different polyps (Figure S1D).

**Figure 2:**
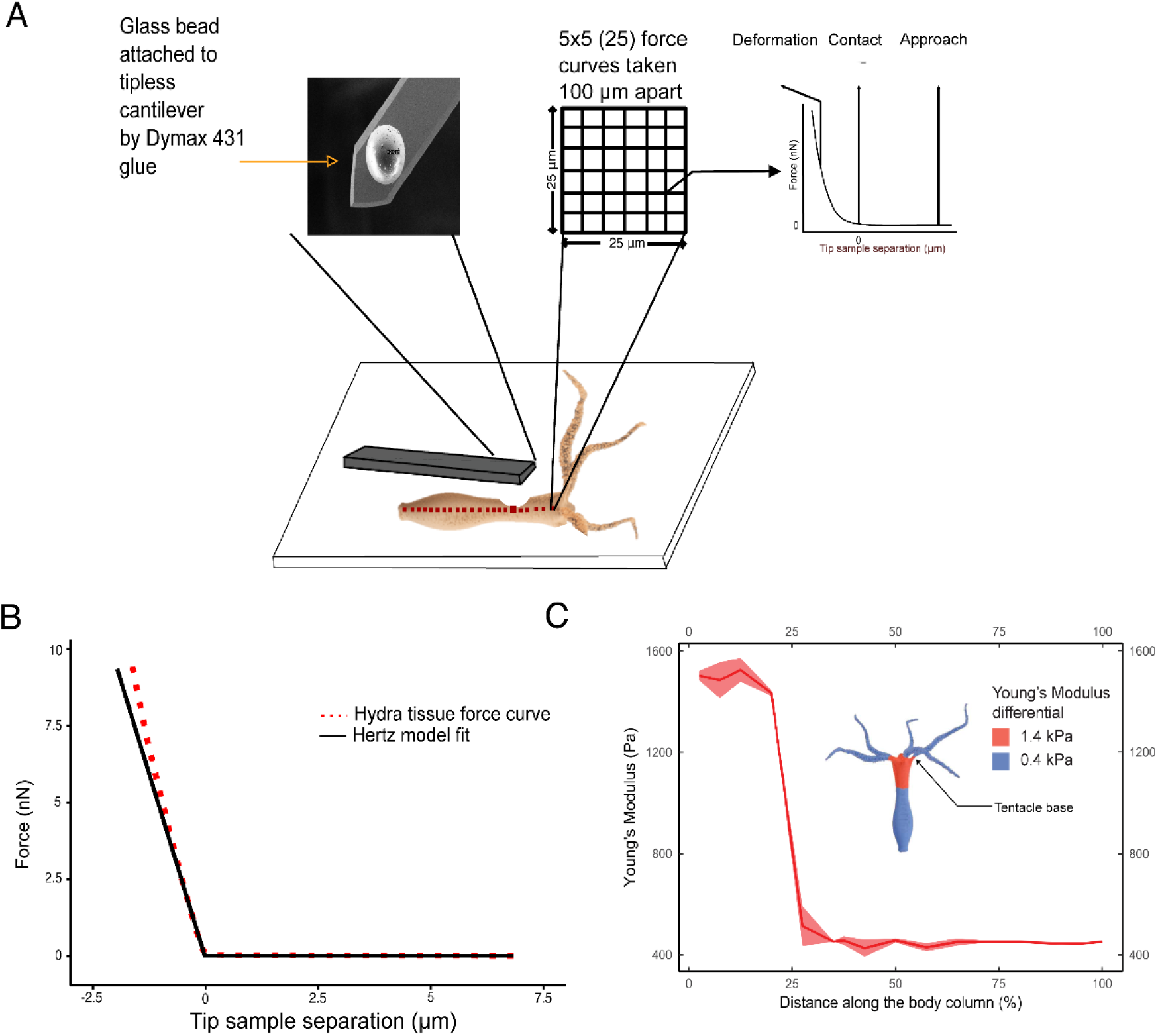
Spatially resolved measurement of Young’s modulus by force spectroscopy. **A.** The schematic of Young’s modulus measurements along the *Hydra’s* body column using an atomic force microscope. A glass bead of 10 μm radius is attached to a tipless AFM cantilever which allows the measurement of the stiffness over multiple locations on the body column. The *Hydra* is attached to glass coverslip using BSA and glutaraldehyde and measurements are taken along the body column over grids separated by 100 μm. Each grid is 25 μm x 25 μm with 25 force curves. Inset shows the image of a bead attached to a cantilever. **B.** A typical force-distance curve used to fit the Hertz model to determine Young’s modulus of a microcontact. The experimental data is shown in red dots, whereas the fit is depicted as a continuous black line. **C.** The plot of variation in Young’s modulus along the body column using AFM equipped with bead attached cantilever. The distance from the tentacle end is plotted in units of percentage of total length. It is zero percent near the tentacles and 100 percent at the base. Force curves were taken at locations separated by 100 μm along the body column for 3 different polyps. The ribbon indicates the standard deviation of the mean over 25 measurements at each location. The cartoon of *Hydra* shows a schematic representation of variations in Y in different regions of *Hydra*. The first (top) quarter of the body column is 3 times stiffer than the rest.

In *Hydra*, the novel sharp change in tissue elasticity along the body column leads to mechanically distinct behavior. The deformations under similar forces would be larger in the body column than the shoulder region and the restorative forces lower in the body column. The observation that *Hydra*’*s* shoulder is three times stiffer suggests that it allows the shoulder to store larger mechanical energy for a given bend. The shoulder region can be viewed as a stiff spring with a high bending rigidity due to its higher Young’s modulus (Figure 3A). This is interesting in context of the somersault, as initially the deformation is seen in the body column and the shoulder region is deformed in the later stages of the movement (Figure 1). The forces acting on *Hydra* to retard its movement in water are mainly viscous in nature. *Hydra*’s weight is mostly negated by the buoyant forces, leading to a reduced gravity situation working against the uplift of the body column. We reasoned that to overcome the viscous and reduced gravitational force on the body column in order to stand upside down, the *Hydra* utilizes energy stored in the bent shoulder. In Fig. 3B, we showed a hypothetical force diagram to better illustrate the forces acting on the organism as the somersualt occurs. For the description we separated the somersault into two phases with the first phase extending the body and stretching the stiff neck region, in the second phase the neck is contracted and bent. The deformation in the neck generates a force F_bend_, which is used to overcome the forces of gravity (F_g_), buoyancy (F_B_) and viscous drag (F_D_).

**Figure 3:**
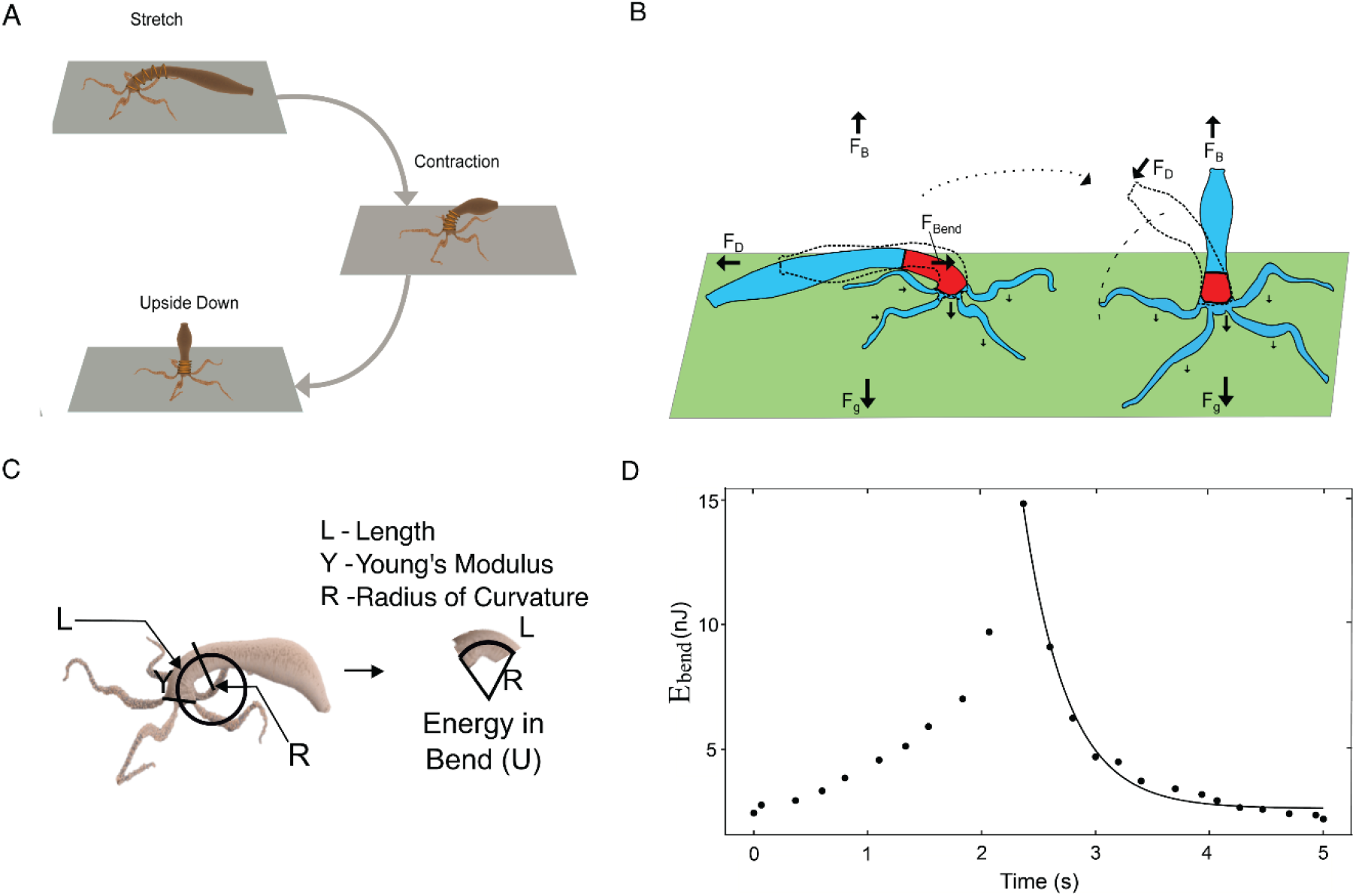
Role of differential tissue stiffness in biomechanics in *Hydra* somersault. **A.** The stiffer shoulder region is depicted as a hypothetical spring. The *Hydra* and the deformations in this spring are shown as *Hydra* somersaults to reach the upside-down position. **B.** The forces acting on Hydra body column during somersault. F_B_ represents the force of buoyancy in water acting against the F_G_ - the gravitational force on the organism. F_D_ is the drag force acting against the direction of motion. F_Bend_ is the representation of the force acting on the head region due to energy stored in the bend. The changes in forces as the *Hydra* goes from Stage 1/Stage 2 (dotted outline) to Stage 4 (dotted outline)/Stage 5 can be seen. **C.** A schematic to illustrate the calculation of energy stored in the bent shoulder. The shoulder region of the *Hydra* is fitted to a circle to measure the radius of curvature (R) of the bend. The length (L) is 25% of the total length (the stiff region), and *Y* is the Young’s modulus as measured by AFM measurements. *I* is the second moment of inertia. **D.** The progression of energy in bend (E_bend_) with time, after the release (t= 0). It first increases to a peak and then exponentially decays as the bend straightens to bring the body to an upside-down position. The continuous line represents a fit to a single exponent (τ~ 0.4 s, n= 1).

Although somersault is an active movement, the observed variation in elasticity points to a possible mechanism to store and release of mechanical energy, which is adopted by animal movement to minimize metabolic costs (Roberts, 2016). Careful analysis of somersault reveals that after releasing from basal end, the bend in the shoulder becomes pronounced. To overcome the viscous and gravitational force on the body column while standing upside down, the energy stored in the bent shoulder is utilized. We analyzed each frame of somersault to estimate the stored energy in the bend (E_bend_) and its progression in time after release from the basal end. The radii of circles measured at different times were used to calculate the bending energy using 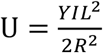, where 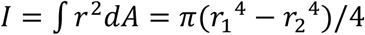 and *U* is the bending energy, *L* is the length of the shoulder region in *Hydra* (25% of body column), *Y* is the Young’s modulus of the shoulder region (stiffer spring), *R* is the radius of the circle fitted to the shoulder region, *I* is the second area moment of inertia, *r* is the perpendicular distance of an elemental area *dA* along the axis of bending, *r_1_, r_2_* are the outer and inner radii of the *Hydra* body column respectively. The change in energy was observed from the point of detachment from the substrate to the final vertical native state of the *Hydra*. We computed E_bend_ using Young’s modulus measured using AFM and radius of curvature R obtained by fitting circles to shoulder region in each frame of the captured video as seen in Figure 3C. Figure 3D depicts the progression of energy in the bent shoulder, E_bend_ with time. The E_bend_ initially increased with time, suggesting a transfer of energy from stretch to the bend. This indicated that after the release, transferring part of the initial mechanical energy into bend might serve as a crucial step in the biomechanics of *Hydra* somersault. *Hydra* lacking a mechanism to facilitate such transfer to optimize the peak in E_bend_ may not have sufficient energy to work against gravity and complete the somersault. After reaching the peak, E_bend_ is used to work against gravity and viscous resistance to reach upside-down position and is expended at an exponential rate (Figure 3D). A single exponent is characteristic of a bent elastic beam relaxing its stress in a viscous environment (Barnes, 1989) and indicates that the motion after release is predominantly governed by passive mechanics. The hypothesis that part of the somersault in which the body is lifted upside down is passive, is further investigated using computer simulations and biological experiments.

### Simulations predict the importance of tissue stiffness variation in somersault

To gain insights into the role of the observed variation in tissue stiffness in facilitating an efficient transfer of mechanical energy to bend, we simulated the part of the somersault which is passive and is depicted in Figure 1. We modelled the tubular body of *Hydra* by a suitable network of 50 rings, each with 10 beads. These rings were stacked together to form an elastic cylinder of length L, as shown in Figure 4A. The individual springs have spring constant *k*. The effective spring constant of the cylinder is *k_eff_*. We observed that *k_eff_* varies linearly with *k* (Figure S2 A and B). The experimentally measured mass, length and Young’s modulus of *Hydra* were used to convert simulation units of m, L and *k_eff_* into physical units (see Appendix 1). To model the experimentally observed stiffness variation along the *Hydra* body-column, spring-constants *k* of individual springs in the shoulder region were chosen to be α times the remainder of the body column (Figure 4A). In simulations, for the same amount of initial energy when the cylinder is stretched (30 nJ), α was varied from 1, which indicates uniform stiffness to a relatively large value of 30 to investigate the importance of spatial variation in tissue stiffness for the mechanics of somersault. Additionally, to truly represent the physical environment, we included viscous drag, the force of gravity and buoyancy on each bead of the model *Hydra* (Figure S3).

**Figure 4:**
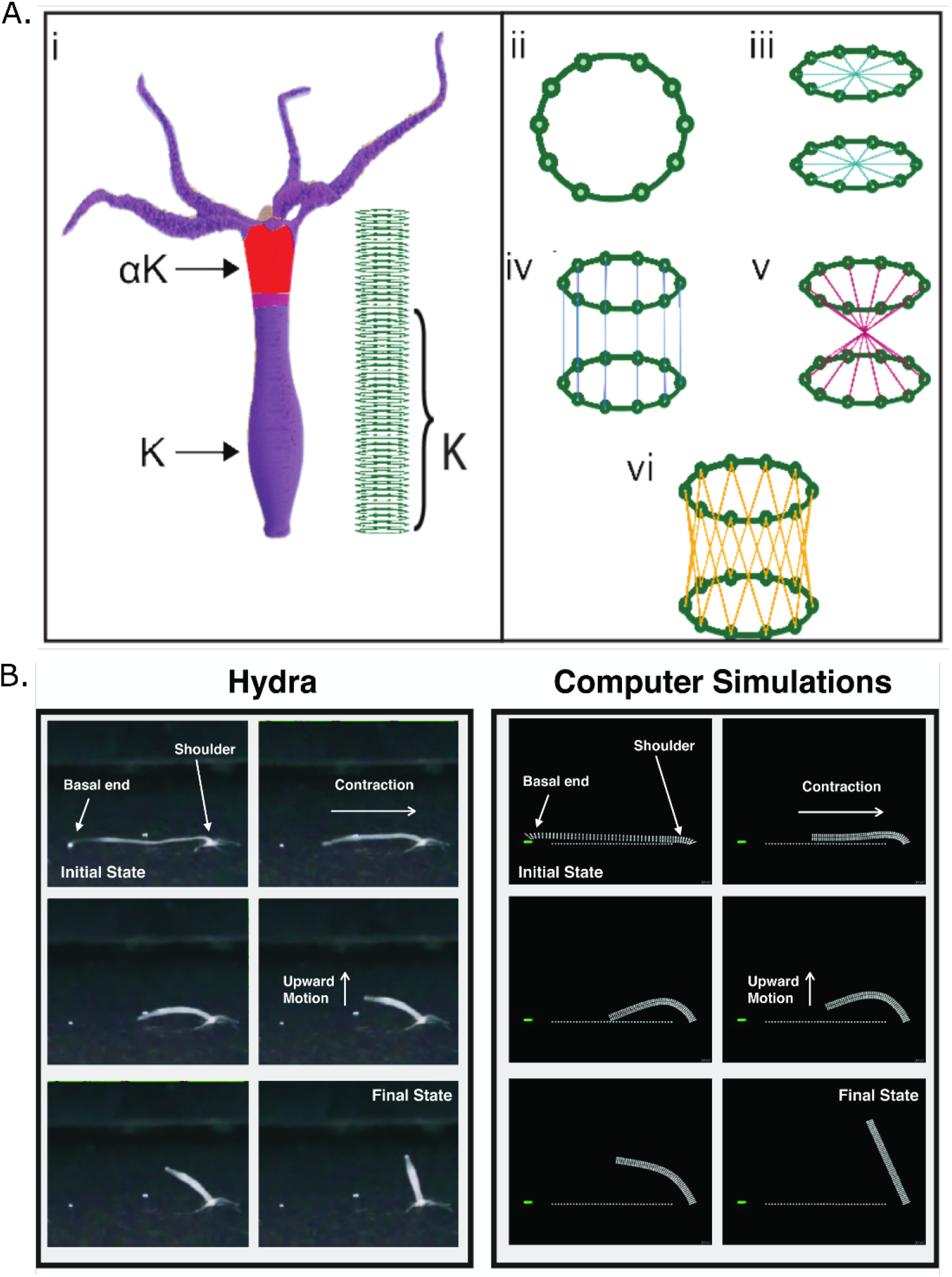
Computer simulations can reproduce the somersault. **A.** Modelling *Hydra* body column to represent various steps involved in the somersault movement. **i)** An elastic cylinder comprising of bead-springs is used to model the *Hydra* body column. The cylinder is a stack of 50 rings, each consisting of 10 beads (mass = m). Beads within and from adjacent rings are connected to each other with springs as shown (ii to vi) to maintain circular cross-section and resist bending, stretching, torsion and shear. The effective spring constant *k_eff_* of a quarter of the body length (shoulder) is kept α times the rest. **B.** The comparison of experimentally recorded various stages of the *Hydra* somersault with simulations. The simulations are performed by incorporating the experimentally observed variation in tissue stiffness. There are striking similarities between both the actual *Hydra* movement and its simulations.

To model the role of elasticity in energy transfer from stretch to bend (Figure 3D), we stretched the cylinder and fixed the two rings at the ends nearly horizontally with strain ε = 0.8 and 0.2. This range of strain values is typically seen in somersault videos. We then released the basal end, which has a lower stiffness. Qualitatively, the somersault in real *Hydra* compares well with simulations for α = 3, which is observed in AFM experiments (See Figure 4B). Interestingly, under uniform stiffness along the length of the cylinder (α = 1), the model *Hydra* was unable to stand upside down after the release (Figure 5A, Movie 2), suggesting that the observed variation in tissue stiffness is critical to complete the somersault.

**Figure 5:**
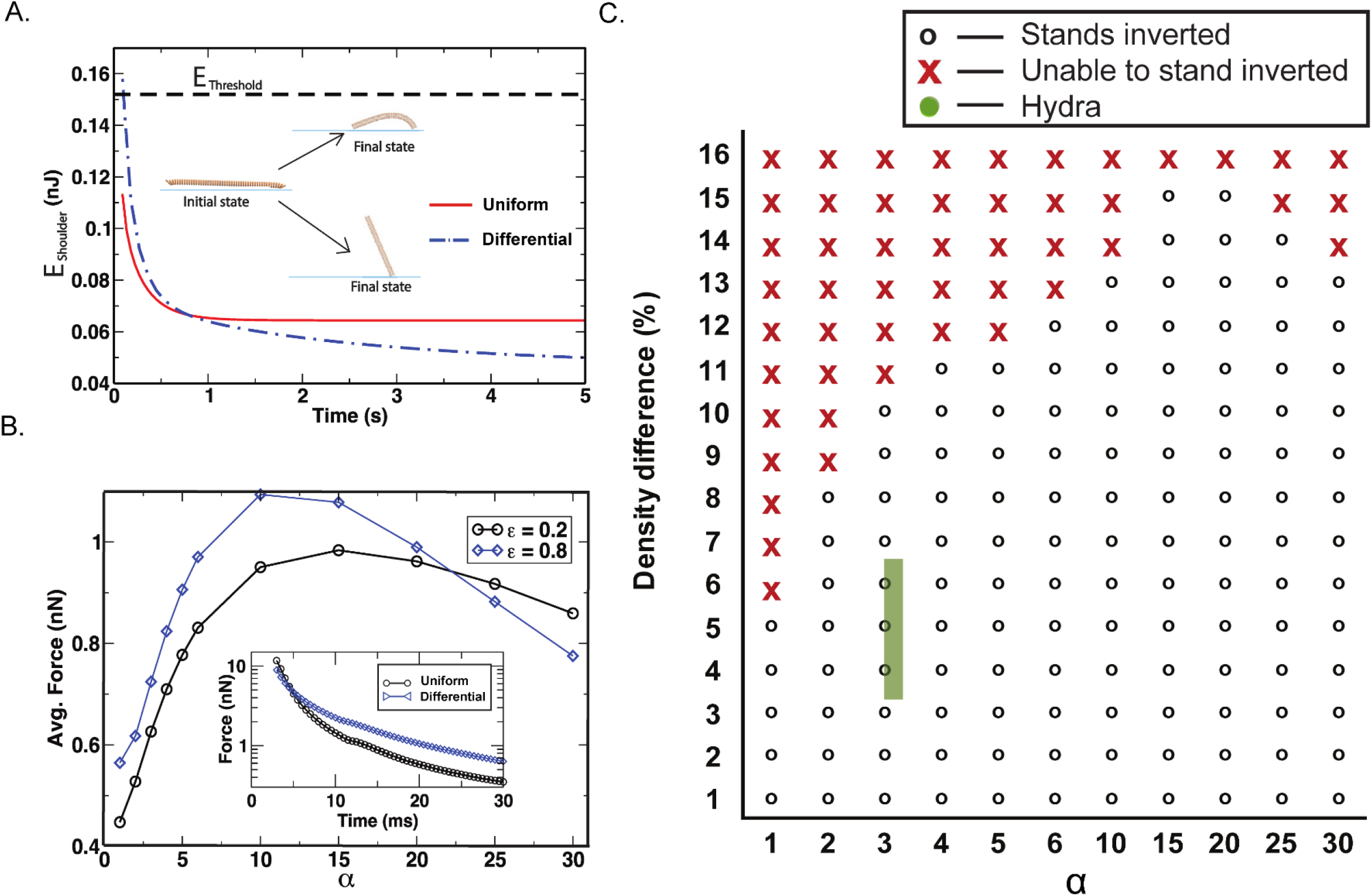
Computer simulations unravel significance of the differential in tissue stiffness for somersault. **A.** For strain ε = 0.8, the plot of the energy in the shoulder region E_shoulder_ versus time after the end of a contraction. E_threshold_ is the calculated minimum energy required to overcome gravity and viscous drag. At the end of a contraction, the E_shoulder for α = 3 is higher than E_threshold and less for α = 1. The simulation snapshots of the cylinder show its initial and final positions for α = 1 (Uniform stiffness) and α = 3 (stiffness differential seen in AFM experiments). **B.** The plot of time-averaged longitudinal force on 14^th^ ring, which is at the junction of the stiff shoulder and rest of the body column, with respect to α after the release. The average is taken over 50 ms before and after the end of a contraction. The force on this ring peaks at an intermediate value of α and does not increase monotonically. This indicates that arbitrarily large values of α do not facilitate the optimal energy transfer. The inset shows force versus time for α =3 and α =1 before the contraction is complete at ~ 50 −100 ms. Initially, the force is nearly the same in both cases, but it becomes roughly twice for α = 3 at later times. Before the contraction is complete, the force is considerably larger for α =3 compared to α=1. **C.** The phase diagram to describe the importance of tissue stiffness variation along the body column to overcome downward force on it due to a higher density of *Hydra* tissue compared to water. The region represented by crosses is the range of parameters in which model *Hydra* is unable to stand inverted after the release, and open circles represent the range in which it is able to stand inverted. The experimentally measured parameters of *Hydra* lie in the green rectangle in the phase space. The width and height of the rectangle represent experimental errors involved in estimating the α and mass density of *Hydra*, respectively. The initial strain ε is 0.8.

The simulations indicated that before the release of the basal end, the potential energy stored in the shoulder for α = 3 was lower than for α = 1. This was due to the comparatively larger stretch in the shoulder for α = 1. However, after release, roughly 50 percent more energy was transferred to the shoulder region in case of α = 3 compared to α = 1. The end of the horizontal movement of the base signifies the end of the contraction. The stored energy in the shoulder region at the end of the contraction step is expended to overcome gravity and viscous drag. The energy threshold (E_threshold_) required to stand upside down can be calculated (Figure 5A, Appendix 1). For α = 1, the stored energy in the shoulder was below this threshold, and for α = 3 it was above it. This suggested that if *Hydra* body column were to harbor uniform stiffness, then it would not be able to stand up (Figure 5A). This result was robust with respect to different values of ε (Figure S4 A and B). It is important to note here that our model treated stage 4 of somersault to be passive. We were encouraged to model *Hydra*’*s* body column as a network of springs and beads due to our observation in Figure 3B wherein, the energy in the bend is expended in a passive manner to work against viscous and gravitational forces. Here, we have focused on the energy transfer process and its utilization to stand upside down. The actual somersault, in its entirety, is a mixture of both active and passive processes and we simulated the passive stage alone. The broad inference that can be drawn from the simulations is that the differential stiffness aids in standing upside down. In particular, it also suggested that the passive energy transfer facilitated by the differential tissue stiffness was a necessary condition for somersault and *Hydra* with uniform stiffness should not be able rise upside down. This prediction from the simulations was then tested experimentally in the next section.

To explain the physics behind the efficient energy transfer for α = 3, we analyzed the simulation data. During contraction, the velocity difference between the shoulder and body column was larger for a model *Hydra* with non-uniform stiffness (α = 3) compared to uniform case (α = 1). This generated more force on the shoulder region while the body column contracts. We calculated time averaged longitudinal force on 14^th^ ring, which is residing at the junction between the stiff shoulder and the labile body column (Figure 5B). Clearly, the force was higher for α = 3 compared to α = 1. The bend in the shoulder, in turn, received adequate energy for standing upside-down. To find out the optimal variation of tissue stiffness for highest possible transfer, we performed simulations for all values of α. Strikingly, the force did not increase monotonically with α but had a clear peak. This indicated that a specific variation in tissue stiffness characterized by α was optimal for energy transfer in *Hydra* somersault and arbitrarily large values of α do not ensure effective energy transfer. In short, the simulations indicated that the variation in tissue stiffness observed upon measurements using AFM is critical for the somersault.

The simulations clearly showed that the variation in tissue stiffness, characterized by α, facilitated efficient energy transfer from a stretch to a bend, which was used to overcome the downward force on the body column due to *Hydra*’*s* higher density compared to that of water. Since this energy was used for overcoming the hydrodynamic drag and the weight of body column due to the density difference (Δρ), the successful completion of somersault depends on Δρ and α. The experimentally measured mass density of *Hydra* is 5.0 ±1.5 percent higher than the density of water (Appendix 1c). We generated phase diagrams of Δρ versus α for range of Young’s modulus values. We observed that for extremely labile bodies (Y < 10 Pa), the energy transferred from the stretched state to the bend is lower than the threshold and *Hydra* is unable to rise for all values of α (Figure S5D). On the other hand, for extremely stiff bodies (Y > 10 kPa), the energy required to stretch to 80 percent strain was large and the transferred energy was always above the threshold such that *Hydra* is always able to rise for all values of α (Figure S5C). These two extremes are far from the experimentally observed stiffness and its variation in real *Hydra* tissue. Figure 5C shows a phase diagram of Δρ and α at the experimentally observed values of Y. Note that in order to keep the amount of energy and strain fixed when the model *Hydra* was stretched, for α < 3, Y for the body region is kept larger than 500 Pa. For α > 3, the Y for the body region was kept smaller than 500 Pa. For Y = 500 Pa throughout the body column, it was not able to rise and stands inverted if the tissue was 2.5 percent denser than water. The experimental measures of *Hydra*, the Δρ and α lie inside the green rectangular shape (Figure 5C). The phase diagram was robust with respect to strain and different initial energies stored in the stretch (Figure S5 A-B). It clearly indicated that a model *Hydra* with larger α, signifying a better energy transfer, can lift itself even if its heavy, which was described by a larger Δρ. It was also interesting that the critical behaviour for lifting the body column was seen in the phase plot of Δρ and α only for the range of tissue stiffness observed in real *Hydra*.

The experimentally measured stiffness of *Hydra* tissue is likely to be an overestimate due to treatment of glutaraldehyde. It was difficult to quantify such effect in case of *Hydra*, however the observed increase in Young’s modulus due to such treatment on rat-tail tendons quantified using AFM is nearly 50 percent (Hansen et al., 2009). We performed simulation runs to monitor the effect of such overestimates on critical nature of standing upside down. Figure S5 A - B shows a phase diagram in which, Y for both shoulder and body column was halved compared to the experimentally observed values in our AFM measurements. The variation characterized by α was seen to be more critical than that depicted in Fig. 5C. *Hydra* with uniform stiffness was unable to lift its body column, even if it is 1 percent above the density of water.

### Modulation of extracellular matrix polymerization perturbs the tissue stiffness variation, impairing the somersault

To identify the source of tissue stiffness and its differential, we hypothesized two possibilities, i.e. variation in the cell types or differences in the extracellular matrix composition. A recent report suggested that there is no difference in cellular stiffness in *Hydra* cells and hence we ruled out the possibility of cellular contribution (Carter et al., 2016). Stiffness variation due to extracellular matrix composition could be tested by selectively disrupting physical properties of mesoglea without disrupting the chemical composition avoiding the possibility of disturbing Integrin-extracellular interactions. *Hydra* mesoglea (extracellular matrix) is a trilaminar porous structure which divides the two germ layers throughout the polyp and is in a perpetually stretched state (Sarras Jr, 2012). Therefore, any lesion to the mesoglea leads to retraction of the mesoglea from the site of injury (Sarras Jr, 2012; Shimizu et al., 2002). The outermost layer of the extracellular matrix is similar to the basement membranes consisting of cell-interacting extracellular matrix proteins such as collagen type IV and laminins (Sarras Jr, 2012). The layer being sandwiched by the basement membrane called the interstitial matrix is primarily made up of fibrillar collagens such as collagen type I, II, III, IV and hence can provide mechanical support to the polyp (Sarras Jr, 2012).

Based on these properties we devised physical and chemical perturbations to mitigate the stiffness of the extracellular matrix. In the physical method, we exploited the retractile property of the extracellular matrix to locally disrupt stiffness which would sever the hypothetical ‘spring’ (Figure 6A). Whereas the chemical method uses Dipyridyl, a lysyl oxidase inhibitor, to inhibit polymerization of the newly secreted fibrillar collagen into fibrils in the extracellular matrix and therefore globally affecting the physical property of newly synthesized extracellular matrix as it replaces old without changing its composition (Figure 6B) (Shimizu et al., 2002; Sarras Jr et al., 1991; Siegel et al., 1970).

**Figure 6:**
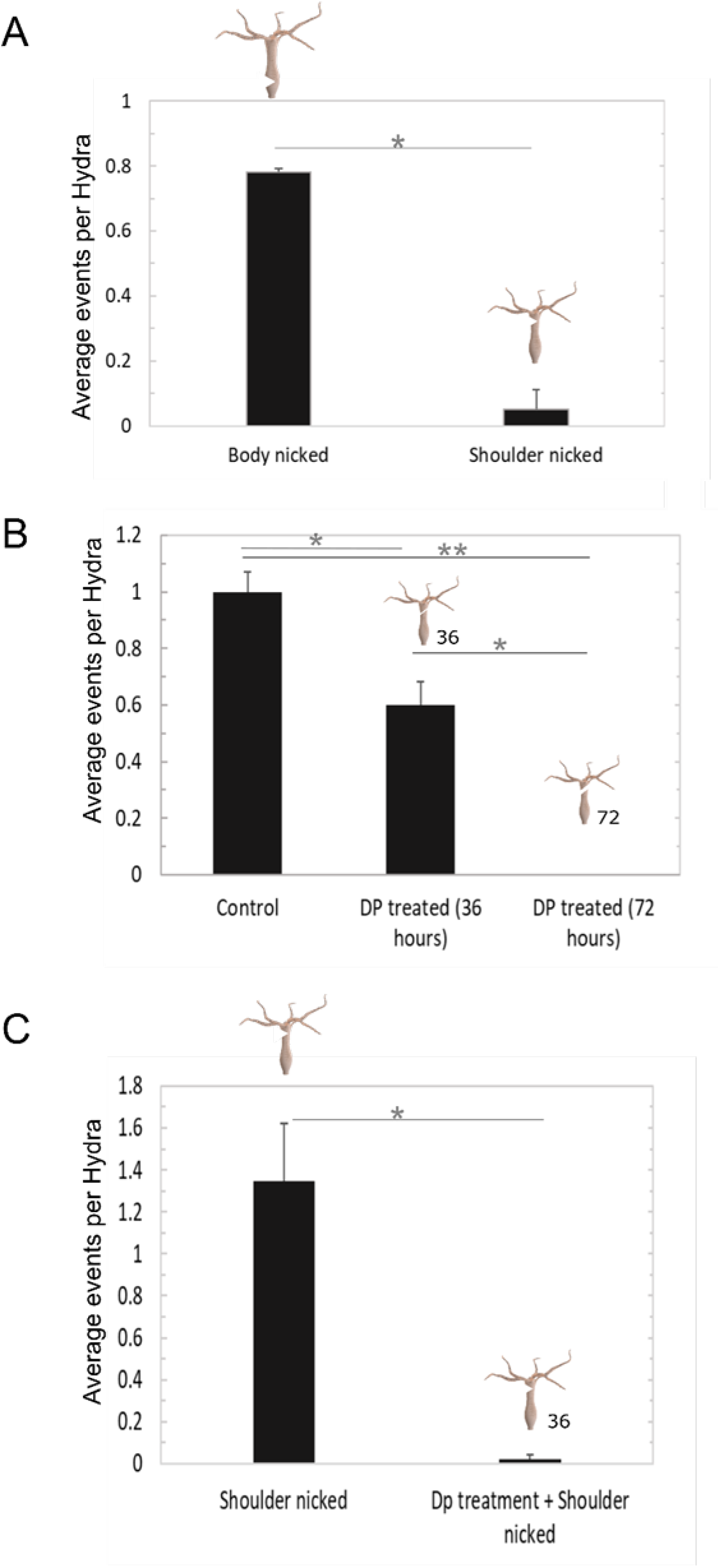
The stiffness differential in the body column is essential for locomotion through the somersault. **A.** The extracellular matrix was perturbed locally in *Hydra* polyp using a partial cut (nick), and the stiffness differential was abolished. The graph shows the average somersault events per *Hydra* with nicks at the shoulder or the body column (number of organisms per experiment =20, number of experiments =3). Perturbing the stiffness in the shoulder region significantly reduces the somersaults by *Hydra* polyps. The error bars represent standard errors, and the significance values are calculated using the 2-tailed Student’s t-test. P-value <0.05 is shown as *, <0.005 is shown as **. **B.** The extracellular matrix was perturbed globally using the chemical disruption of collagens by treatment with 10 mM Dipyridyl (number of organisms per experiment =20, number of experiment =3). The average somersault events per *Hydra* reduce upon treatment with Dipyridyl for 36 hours, and none are observed after treatment with Dipyridyl for 72 hours. The error bars represent standard errors, and the significance values are calculated using the 2-tailed Student’s t-test. P-value <0.05 is shown as *, <0.005 is shown as **. **C.** The differential in stiffness was perturbed by disrupting the extracellular matrix with a combination of Dipyridyl treatment and a partial nick. As shown in the graph, abolishing the stiffness differential with a combination of physical nick and Dipyridyl treatment also leads to a reduction in the average number of somersault events observed. The animals are nicked in the shoulder and treated with Dipyridyl for 36 hours. The error bars represent standard errors, and the significance values are calculated using the 2-tailed Student’s t-test. P-value <0.05 is shown as *, <0.005 is shown as ** (number of organisms per experiment =20, number of experiment =3).

On two different sets of *Hydra* polyps, we partially lesioned the shoulder region and the middle of the body column severing the extracellular matrix (number of polyps per experiment =20, number of experiments =3). Then we recorded the somersaults performed after 6 hours after lesion. This time point was chosen since it is sufficient to heal the wound but not enough for regaining the stiffness around the lesion because the extracellular matrix is not fully regenerated(Sarras Jr et al., 1991, Shimizu et al., 2002) (Figure 6A, movie 3). Further, the stiffness measurements using AFM revealed the loss of differential stiffness seen in the *Hydra* body column (Figure S6 A). Strikingly, the ability to perform somersaults was virtually lost when the lesion is affected in the shoulder region, whereas it remains unaffected when the lesion was in the body column (Figure 6A). The polyps with a lesion in the shoulder were able to regain the ability to somersault after 36 hours, a duration long enough to complete the regeneration of the extracellular matrix. Thus, elimination of the stiffness locally within the shoulder region resulted in an inability to perform the somersaults, underscoring the importance of the stiffness differential. The stiffness measurements using AFM revealed that the polyps treated with Dipyridyl for 72 hours exhibit uniform stiffness across the entire body column (Figure S6 B). This suggested that *Hydra* presumably exploits the differential crosslinking of fibrillar collagen to generate tissues with varying degree of elasticity. The ability to somersault was completely lost after 72 hours of Dipyridyl treatment, whereas the number of somersaults was reduced to half after 36 hours of treatment (Figure 6B and Movie 4) (number of polyps per experiment =20, number of experiments =3). This confirmed the correlation of the degree of unpolymerized fibrillar collagen in the extracellular matrix to the inability to somersault. These observations suggest that the extracellular matrix mediated stiffness differential in the *Hydra* polyp is critical for somersaulting. In the third experiment, the animals with a lesion in the shoulder region were treated with Dipyridyl. This does not allow the regeneration of the extracellular matrix in the nicked region. In this case, even 36 hours after the nicking the polyps were unable to regain the propensity to somersault (Figure 6C and Movie 5).

To probe deeper into the role of tissue stiffness as a function of collagen crosslinking, we monitored the ultrastructure of the extracellular matrix in mesoglea using scanning electron microscopy (SEM). Tissue sections were prepared from the shoulder region and body column to reveal the difference, if any, in the ultrastructure of the extracellular matrix reflecting the tissue stiffness differential. SEM images of these sections revealed clear demarcation of the mesoglea (me), indicated in the orange dotted line separating the ectodermal tissue (ec) from endodermal tissue (en) (Figure 7). Within mesoglea, due to the transverse sectioning of the tissue, the central interstitial matrix consisting of fibrillar collagens were distinctly visible (Figure 7 A’, B’, C’ & D’). In the control polyps, comparison of the SEM images clearly depicted the differential packing of the extracellular matrix collagen fibers and indicated that this could be attributed to differential collagen crosslinking (Figure 7 A-A’ & B-B’). These observations suggested that the stiffness differential observed between shoulder region and body column was presumably due to the differential packing of collagen fibers in the extracellular matrix. As discussed above, Dipyridyl treatment of *Hydra* polyps resulted in the loss of differential stiffness between the shoulder and the body column by inhibiting crosslinking of the collagen fibers (Figure S6 B) at 72 hrs. To evaluate such an effect on the ultrastructure of *Hydra* extracellular matrix, we monitored the extracellular matrix structure in polyps after Dipyridyl treatment. We performed SEM imaging of tissue sections of Dipyridyl treated polyps in the shoulder region and body column. The polyps treated with Dipyridyl for 72 hrs corroborated the results obtained through stiffness measurements using AFM (Figure 7 C-C’ & D-D’). Inhibition of lysyl oxidase activity by Dipyridyl caused an extensive reduction in collagen crosslinking in the shoulder region (Figure 7 C-C’) as compared to the control polyps. The extracellular matrix ultrastructure at the body column after Dipyridyl treatment appeared unaffected relative to the control polyps. This observation also confirmed that collagen in the mesoglea of the body column is sparsely crosslinked as compared to the shoulder region. Collectively, these findings provide a compelling evidence that spatial variation in tissue stiffness is a function of collagen crosslinking and is critical for the somersault, corroborating the results of the simulations.

**Figure 7:**
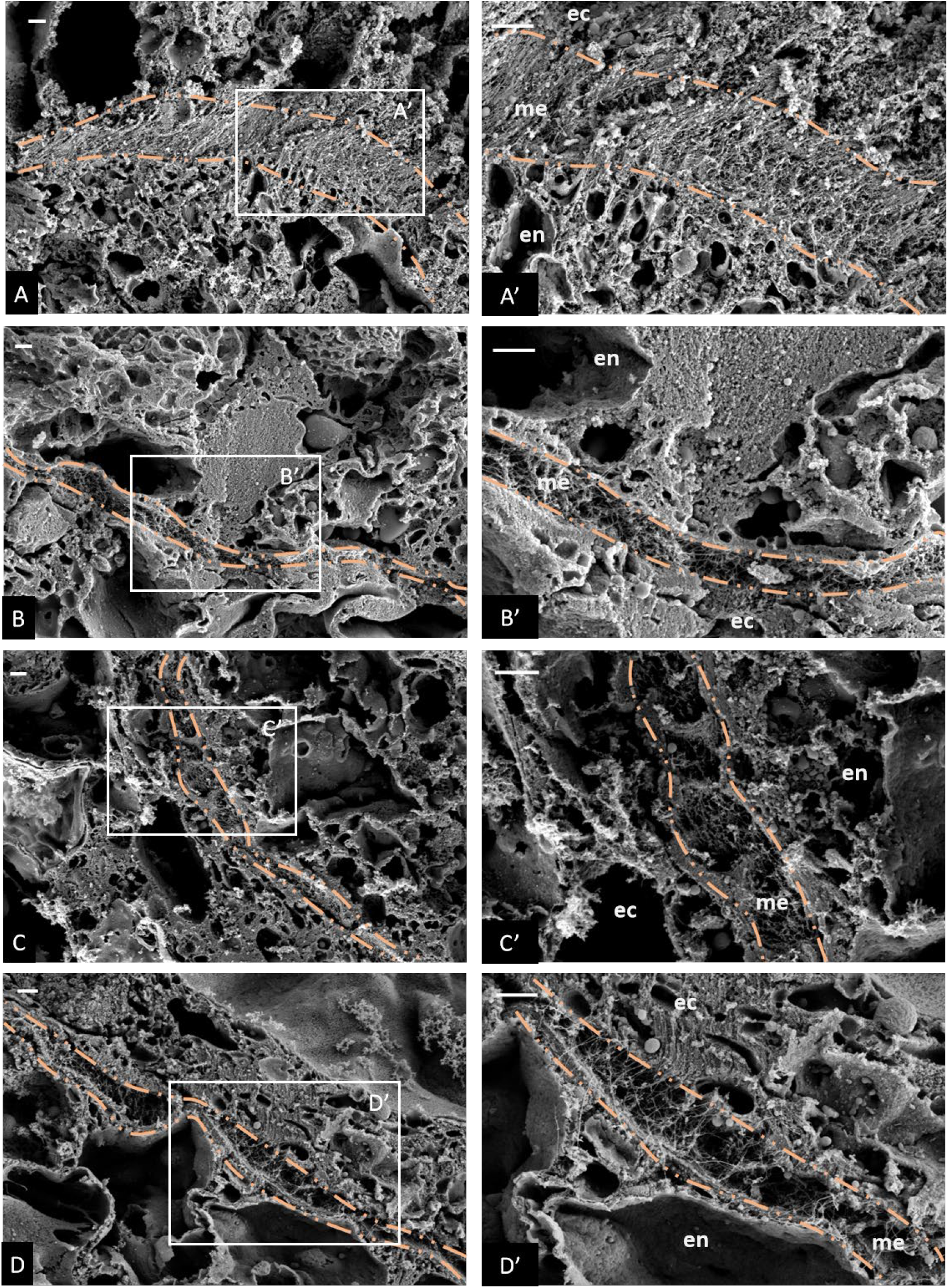
Scanning electron micrographs (SEM) of *Hydra* mesoglea showing changes in extracellular matrix between shoulder region and body column in control (A-B) and Dipyridyl treated (C-D) polyp. **A.** SEM image of mesoglea (orange dotted region) from the shoulder region of the control polyp in transverse section (TS) showing dense collagen fibres. A’ shows higher magnification image of region indicated by a rectangle from A. Scale bar= 1 μm (both for A and A’). **B.** SEM image of mesoglea (orange dotted region) from the body column region of the control polyp in TS showing less dense collagen fibres. B’ shows higher magnification image of region indicated by a rectangle from B. Scale bar = 1 μm (both for B and B’). **C.** SEM image of mesoglea (orange dotted region) from the shoulder region of the polyp after Dipyridyl treatment for 72 hrs in TS showing loosely packed collagen fibres due to inhibition of collagen crosslinking. C’ shows higher magnification image of region indicated by a rectangle from C. Scale bar = 1 μm (both for C and C’). **D.** SEM image of mesoglea (orange dotted region) from the body column region of the polyp after Dipyridyl treatment for 72 hrs in TS showing loosely packed collagen fibres due to inhibition of collagen crosslinking. D’ shows higher magnification image of region indicated by a rectangle from D. Scale bar = 1 μm (both for D and D’). Abbreviation: ec- ectoderm, ec- ectoderm & me- mesoglea.

## Discussion

We report the first-ever use of atomic force microscopy (AFM) for measuring the tissue stiffness of *Hydra* along the whole body-column. We have obtained more definitive results than the earlier reports of tissue stiffness in *Hydra* (Carter et al., 2016). Here, we demonstrated that *Hydra* possesses a local variation in tissue stiffness at a ratio of 3:1 between the shoulder region to the rest of the body column.

To understand the significance of such a sharp stiffness differential in the body column, we investigated the seemingly strenuous process of somersaulting (Biewener, 1990). The simulations revealed that the observed differential in the tissue stiffness is necessary for energy transfer to stand upside down. In congruence with simulations, the *Hydra* polyps lacking the stiffness differential were unable to stand upside down during somersault. The differential speeds of shoulder and rest of the body during retraction leads to a soft collision between the two regions, which results in an efficient transfer of energy from the body column to the stiff region (shoulder). Further, the differential stiffness of the body column enables the animal to strike an optimal balance between the mechanical energy stored in the stretching and the bending processes. While stretched, most of the energy is stored in the labile body column due to the extension. The collision allows for the energy produced in the extension step to be concentrated into potential energy in the bend of the shoulder region. Comparing the somersault to walking on land, the arrangement of the shoulder and the body column acts similar to the movement of legs. The tentacles allow for stability by providing lateral forces similar to insect legs (Dickinson et al., 2000). The shoulder region acts like a spring storing energy from the stretched body column and repurposing it to lift vertically. While somersaulting allows the polyp to move around in the immediate environment in intermediate distances, looping allows the polyps to cover smaller distances. For larger distance locomotion, floating using an air bubble seems more favorable. Whereas, the labile part of the body column can extend with relative ease allowing the *Hydra* to probe its immediate surroundings without moving to a new position and expending too much energy. We also observed in our experiments that the polyps showed a higher propensity to perform looping, somersaulting or stretching when we detached them and dropped them back in the culture bowl. This behavior can be explained by the need for *Hydra* to probe the immediate surrounding to gauge the nature of the new site when it settles in natural conditions. Such probing might be important in obtaining different sets of information by the polyp including water flow differential in the surrounding, better substrate attachment sites, prey abundance etc.

The arrangement of the shoulder and the body column can be compared to that of tendons and muscles in vertebrates. The spring mechanism in the shoulder region is similar to mechanisms proposed for motion during terrestrial walking (Markowitz and Herr, 2016, Taylor and Heglund, 1982). Energy recycling through tendons and other elastic elements is an established mechanism in describing locomotion in vertebrates such as humans (Cavagna et al., 1977). Tendons consist of thick bundles of collagen fibres secreted by tenocytes. Incidentally, tenocytes can change the extracellular matrix composition of tendons in response to changes in mechanical loading and hence the elasticity (Hansen et al., 2009). Based on our study, mechanisms modulating elastic properties of tissues to facilitate locomotion seems to have evolved early in the animal phyla and hence seems to be an ancient function of the extracellular matrix. It points to the possibility that shear stresses faced during tentacle movement as well as the structural peculiarities of the organism may have played a role in evolving the stiffness differential (Carter et al., 2016; Shostak et al., 1965; Haynes et al., 1968). The torsional forces for moving a flexible rod-like structure as tentacle in water could be of the range of nanonewton/micronewton and deformations in the hypostome would be of the order of nanonewton. Such scales of repetitive deformations could have led to the evolution of mechanosensory feedback mechanisms required for regulating the extracellular matrix stiffness. One of the easiest ways to achieve the same is by modulating the crosslinking of fibrillar collagens in the extracellular matrix, an example of which we have illustrated here. A similar mechanism might have further evolved in more complex higher organisms during the development of the musculoskeletal system.

Our study underscores the importance of the observed changes in cellular and molecular properties at the shoulder region and their mechanistic contribution to the process of locomotion in *Hydra*. The cellular movement shows a marked difference between the shoulder region and the body column (Holstein et al., 1991). All the cells in a polyp are known to be pushed towards the termini of the body and sloughed off. The extracellular matrix associated with these cells is also reported to move along with these cells and degraded at the tentacle tip or the basal disk (Aufschnaiter et al., 2011). Strangely, the aforesaid behavior changes at the shoulder region with a reduction in the cell-extracellular matrix velocities. This could be due to differential interactions of cells with the components of the extracellular matrix in the shoulder region versus the rest of the body column, as has been proposed earlier (Shostak et al., 1965). The extracellular matrix is degraded within the shoulder region than being pushed to the tentacles indicating that the extracellular matrix in this region is maintained in a different manner as opposed to other regions. Further, BrdU staining indicates a cessation of the proliferative capacity of the stem cell as they cross the boundary towards the shoulder region and into the hypostome and tentacles (Reddy et al., 2015). In *Hydra*, the tentacles and the hypostome region possess a wide range of differentiated cells, which terminally differentiate as they move from the gastric region towards the hypostome and tentacles (Wood, 1979, Ewer and Fox, 1947). It would be interesting to study if, similar to human and mice cells, stiffness properties of the extracellular matrix determine the differentiation fate of these cells in *Hydra* (Engler et al., 2006, Park et al., 2011). Thus, tissue stiffness in *Hydra* is not only innately linked to the motility of the organism but also possibly other processes.

Collagen, as shown here, plays an important role in the tissue stiffness, possibly both through cell-extracellular matrix interactions and varying extracellular matrix composition (Shostak et al., 1965; Aufschnaiter et al., 2011; Zhang et al., 2007; Shimizu et al., 2008; Sarras Jr, 2012). The collagen fibre structure in the extracellular matrix can be dynamically regulated depending on the strain experienced locally (Haynes et al., 1968). Studies need to be performed to understand how collagen and other extracellular matrix molecules are regulated along the oral-aboral axis of *Hydra*. Our findings not only open a new avenue in invertebrate locomotion mechanics which could lead to interesting insights into the evolution of locomotion but also lay a foundation for furthering mechanobiology in the lower organisms using AFM based measurements. Additionally, the need of an optimal elasticity variation for greater energy efficiency in liquid environments could be a significant design principle for building artificial machines and advance the field of the untethered small-scale robots working in confined areas(Hu et al., 2018).

## Methods and Materials

### 1. *Hydra* lines and culture

The polyps used were the Pune strain of *Hydra vulgaris* (Ind. Pune) and *Hydra vulgaris* (AEP) (Reddy et al., 2011; Martínez et al., 2010). *Hydra* polyps were cultured under standard conditions in glass bowls with *Hydra* medium containing KCl, NaCl, MgSO_4_.7H_2_0, CaCl_2_.7H_2_O and Tris-HCl using the protocol described previously (Lenhoff, 2013). Both lines were maintained at 19°C and with a day night cycle of 12 hours. Polyps were fed using freshly collected Artemia hatched in house. The medium was changed every day after feeding.

### 2. Measurement of *Hydra* tissue stiffness using atomic force microscope (AFM)

#### 1. Attaching bead to the AFM lever

A tipless cantilever with a glass bead attached to its free end (stiffness ~ 0.2 N/m) was used for AFM measurements. The diameter of the glass bead is 20 μm (Figure 2 A). The attachment was accomplished using the micromanipulation available with the AFM. A small amount of UV curable glue (Dymax 431) was spread on the coverslip. Using the servo control of the AFM, the end of the tipless lever was lowered onto the glue. A drop of glue was picked up on the lever and lowered again on a bead. The lever was maintained under positive load and UV light was directed at the bead-lever assembly. After curing the lever was pulled back from the surface along with the bead. The elastic modulus of the glue is 570 MPa, and it is not deformed while pushing on the tissue. Before performing force-distance measurements, the cantilever is calibrated using BSA coated glass surface by thermal tuning.

#### 2. Measurement of tissue stiffness using AFM

*Hydra* body is unstable for mechanical measurements if it is not strongly adhered to the glass surface. A thin layer of BSA was coated on the glass for strong adherence of the tissue. Young’s modulus of various tissues typically ranges from 100 Pa to 1 MPa. The Young’s modulus of glass is of the order of 10-100 GPa. The coating of BSA alters it to some extent. Figure S1 A shows the cantilever deflections for glass, BSA coated glass and tissues from different parts along the body column. Assuming the glass-glass contact to be infinitely stiff compared to the glass-tissue contact - a reasonable assumption since it is 10,000 times stiffer, the slope of the curve in the contact region for glass and BSA coated glass is nearly one implying no deformation. The slope of the curve on tissues is much less, suggesting a certain amount of deformation. We use glass-glass contact for calibration of deflection sensitivity and the subtraction of cantilever deflection from the push given by the piezo extension yields deformation in the tissue. The force is calculated by multiplying the cantilever deflections by its stiffness. The force versus deformation curve is then fitted with the Hertz model.

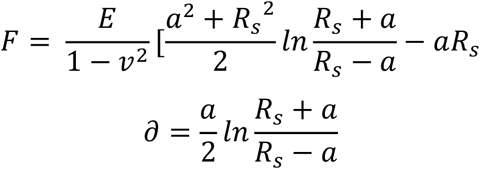

Where F is the measured by the cantilever possessing the bead, which is pressed against the tissue. R is the bead radius, the delta is the deformation in the tissue, E is Young’s modulus, and v is the Poisson ratio.

Hertz contact mechanics theory works for non-adhesive elastic contacts. It is important to establish that the pressing of bead over the tissue conforms to this requirement. Figure S1B shows a typical force-deformation measurement while both extending and retracting the bead over the tissue. For small loads (< 1 nN) the extend and retract curves do not show hysteresis, which indicates that the contact is non-adhesive and elastic.

Before AFM measurements, *Hydra* polyps were cultured and starved for a day to eliminate food material. They were relaxed with urethane (2% for 2 mins) and fixed immediately with glutaraldehyde (4%) for 30 mins. The coverslip (diameter: 22 μm) was coated with a layer of bovine serum albumin (BSA, 10 mg/ml), and this layer was allowed to dry. The *Hydra* was placed on this layer, and a small amount of BSA was added to keep it from drying. As the BSA dried, the connections formed between the *Hydra* and the surface by BSA were fixed using glutaraldehyde for 2 min, and water was added. The errors in the measurement of Young’s modulus determines the width of the green rectangle used to depict the experimental measures of *Hydra*.

### 3. Measurement of the mass density of *Hydra*

The density of *Hydra* was measured by the following experiment. Tentacles were removed and the resulting *Hydra* body column was dropped in the water column in a vertical tube (height 2 m, diameter 5 cm). The body column attains terminal velocity and is measured to be 0.003 m/s. The body column moves downwards horizontally without rotating or tumbling (Movie 6). After balancing the forces acting on a cylindrical body moving with a terminal velocity in a fluid, we obtain

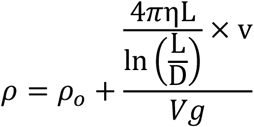

Where ρ is the density of *Hydra*, ρ_0_ and η are density and viscosity of water respectively. L is the length of *Hydra* body column, D is the diameter of the body column and V is the volume of *Hydra*, g is the acceleration due to gravity. We measured the length and diameter at five different locations across the body column of ten different *Hydra* polyps. The errors in the measurement of L and D largely determine errors in estimating the density of *Hydra* from the measurements. The density of *Hydra* tissue is 1050 ±15 kg/m^3^. This is 5±1.5 % above the density of water. This error determines the height of the green rectangle used in Figure 5C.

### 4. Videography and analysis of somersault

We recorded the motion of *Hydra* as it does the somersault movements and measured the statistics of somersaults with modification of native stiffness along body column (Figure 6). We fabricated a glass tank of 10 x 8 x 4.5 cm dimensions and recorded the movement using a Nikon D-500 digital camera fitted with a 105 mm macro lens. The *Hydra* movement was recorded as they detached from the surface. For measurement of distances a precise scale was kept in the bath. The videos were recorded at 25 fps for 20 min at a stretch. We assayed the propensity of *Hydra* to somersault after dropping them with a pipette into the tank. For the ease of scoring, we have only scored the number of times polyps stand upside-down since it is the first and most important step for *Hydra* to somersault. The normalization of the events was done by calculating the average and standardizing the values by comparison to the control. The videos were analyzed with the Light orks software and Fiji. For bending energy calculations in Figure 3D, circles were fit to the shoulder region of *Hydra* using Fiji software and the bend energy stored in the shoulder region was measured.

### 5. Extracellular matrix disruption

*Hydra* extracellular matrix has been shown to be dynamically regulated and important in regeneration as well. It has been seen before that an amputation leads to a retraction of the extracellular matrix near the wound. This retraction is about 100 μm below the cut edge, and the extracellular matrix re-secretion takes about 24 hours to complete (Sarras Jr et al., 1991). We used this property of the extracellular matrix to physically perturb it upon nicking (partial cut/amputation) of the organism. The nicking of *Hydra* is performed using the sharp bevel edge of a 31-gauge syringe needle and carefully controlled under a 10X dissection microscope. Adequate care is taken such that the nick always is halfway through the body (incision up to half of the diameter of the polyp at the shoulder region) and perpendicular to the long body axis, such that the severed part does not move away. It is assured that the head attaches in the same place, and the wound is healed to carry out the videography of the somersaulting locomotion. Such amputation would lead to a loss of extracellular matrix in a small region around the partial amputation. To assess the effect of loss of extracellular matrix in a particular region on the stiffness of the organism, we performed AFM-based measurements on head partially cut polyps. To assess the effect of localized extracellular matrix perturbation on ability of *Hydra* to somersault, the polyps were videographed for somersault 6 hrs post amputation. Twenty polyps per experiment were used and a total of 3 experiments were performed with no polyps reused for any experiments. These polyps were scored for ‘Events per *Hydra*’ The scoring was performed by counting total number of occurrences the polyps performed upside-down movement from the time they were dropped in the container floor for the first 5 mins. These movements are labeled as ‘events’ and these events were then divided by total number of polyps in the container to obtain the value of ‘Events per *Hydra*’ (Figure 6).

2,2’-Dipyridyl (Dipyridyl) is an inhibitor of lysyl oxidase (Siegel et al., 1970). Lysyl oxidase is an enzyme which crosslinks two adjacent fibrillar collagens to make bundles. Inhibition of lysyl oxidase prevents the components of the extracellular matrix from polymerizing. This has been shown before in being effective in inhibition of *Hydra* extracellular matrix [6]. The concentration of 100 μM was shown to be useful in other strains of *Hydra*. To test the validity of this result and to study the behavior of *Hydra* under this drug concentration from 50 μM to 200 μM Dipyridyl in *Hydra* medium were tested, and no physiological effects were detected until 175 μM. For the partial amputation along with Dipyridyl treatment experiments (Figure 6 C), the organisms were pre-treated with Dipyridyl for 12 hours followed by nicking and further treatment with Dipyridyl for 24 hours. *Hydra* extracellular matrix is continuously cycling along the body column and replenished. Twenty polyps were used per experiment, and a total of 3 experiments were performed with no polyps reused for any of the experiments. For the Dipyridyl treatments (Figure 6 B), Dipyridyl was added to the *Hydra* medium at 100 μM, and the medium was changed every 24 hours. Stiffness measurements were performed using AFM on *Hydra* treated for 72 hours to assess the effect of chemical extracellular matrix disruption on the stiffness.

For the comparison between different experimental perturbations to the *Hydra*, we compared 20 animals for each condition in a biological replicate. Each biological replicate for an experimental perturbation was repeated with its respective control for three trials. The average number of somersaults for each condition was measured during a fixed time period. The averages for the replicates were compared with corresponding controls, and significance was measured using the unpaired Student’s t-test.

### 6. Scanning electron microscopy of *Hydra* tissue

Anaesthetized polyps (using urethane (Sigma)) were fixed using EM fixing solution (2.5 % Paraformaldehyde (Electron Microscopy Sciences - EMS), 2.5 % Glutaraldehyde (EMS) and 0.1 M Cacodylate buffer (Sigma)). These polyps were then embedded in 5 % Agar and sectioned into 200 μm slices using a vibratome (Leica VT1200 S). The sections were then stained with 1 % Osmium tetroxide (Sigma), 1% Tannic acid (EMS) and 1 % Uranyl acetate (EMS). These sections were then dehydrated by washing them serially with increasing concentrations of ethanol (30-100 % ethanol). These sections were then dried at CO2 critical point using a critical point dryer (Leica CPD030). These were then mounted on carbon tapes and sputter-coated with platinum. The coated samples were then imaged using the Sigma 500 electron microscope (Zeiss).

## Appendix 1

### Simulation Units of all physical quantities

The experimentally observed values of the length of the *Hydra* body column is around 5 mm while its inner and outer radii are 0.05 mm and 0.1 mm, respectively. In our simulation, this is modelled as a hollow cylindrical tube of diameter of a = 1 simulation units (s.u) and length L0 = 30a (see Table A1).

**Table A 1.**

| Dimension | Simulation Units | SI Units |
| --- | --- | --- |
| Length [L] | $a = 1$ | $0.17 \times 10^{-3} \text{ m}$ |
| Mass [M] | $m_i = 1$ | $2 \times 10^{-10} \text{ kg}$ |
| Time [T] | $\tau = 1$ | $0.91 \times 10^{-3} \text{ s}$ |

The stretched elastic body of *Hydra* can be physically characterized by its Young’s modulus Y, stored elastic energy E, length L, cross-sectional area A (found using its inner and outer radii) and its mass M= ∑mi, whereas viscous drag due to liquid surrounding *Hydra* can be characterized by the viscous drag coefficient Γ. These parameters can be combined to form a dimensionless parameter *D*.

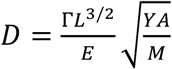

Substituting 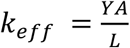, which is essentially replacing the Young’s modulus by the stiffness constant of an equivalent spring, gives us

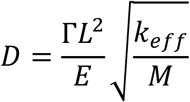

We can estimate these parameters in the following way:

#### a. Elastic energy (E)

We take the experimentally measured value of the Young’s modulus around 1000 Pa, and we assume the stress-strain relationship linear, i.e. σ = Yε. Stored elastic energy per unit volume is given by the integral of the stress-strain curve. For the sake of simplicity, we take energy per unit volume, stored at a strain of ε = 1

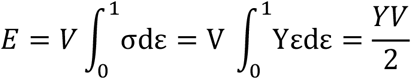

Using the values of inner and outer radii for the *Hydra* as well as its length of 5 mm, we get the energy value of the elastic energy to be

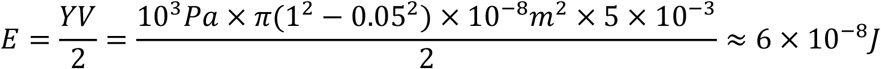

#### b. Stoke’s friction coefficient for a Cylinder (Γ)

The drag force experienced by the *Hydra* body column is determined by its Stoke’s coefficient. The diameter to length ratio D/L of *Hydra* is small fraction, suggesting that it can be approximated as a long thin cylindrical tube (Akhtar et al., 2011; Dhont, 1996). The drag coefficient under this approximation is divided into two parts one parallel to the cylinder axis (running along its length), which we call Γ^∥^, and the other perpendicular to the axis Γ. Ideally, Γ^∥^ acts during the initial contractile motion of the *Hydra*, whereas Γ^∥^ acts during the upward motion. The expressions for these coefficients are

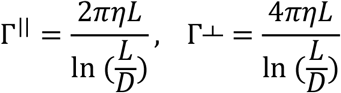

Here, L and D are length and diameter of *Hydra* body, while *η* is the viscosity of water, which is the environment they exist in. It can be seen that Γ^∥^ = Γ^⊥^ 2. Since both these values do not defer by an order of magnitude, we take their average value as the drag coefficient

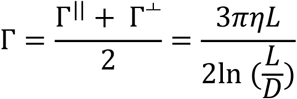

Putting the viscosity of water as 10^−3^ Pa, L = 5 x10^−3^ m and D = 10^−4^ m, we get Γ = 1.2 x 10^−5^ Ns/m

#### c. Mass (M)

We have measured *Hydra’s* body to have a mass density of 1050 kg/m^3^, and its mass can be estimated as *Hydra’s* volume multiplied by its density. This comes out to be about 10^−7^ kg.

#### d. Stiffness Constant of equivalent spring (*k_eff_*)

We use the relation of *k_eff_* to calculate this as

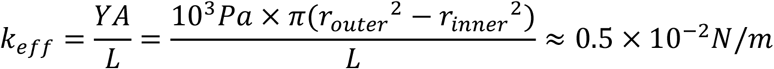

Using these values, the theoretically estimated value of the dimensionless parameter D is calculated as

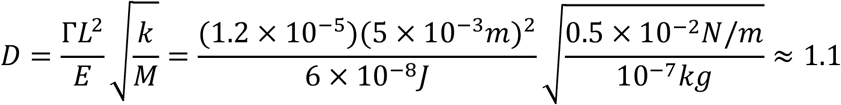

The simulation parameters are chosen such that this dimensionless quantity remains unchanged when calculated using the simulation units. A choice of a stiffness constant fixes the stored elastic energy of the system. Using the effective stiffness value of 20 s.u, the stored energy can be calculated as ½ *k_eff_*. (Δx_sim_)^2^. The neutral length of the simulation *Hydra* being 30 s.u., upon being stretched to twice its length gives the value of Δx_sim_= 30 s.u. Using these values along with the simulation mass as 500 s.u. (each bead being assigned a mass of 1unit, with 500 beads forming the cylinder), we get

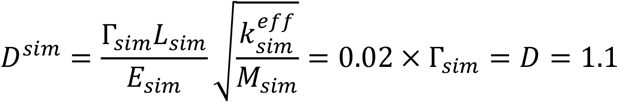

Solving for Γsim, the value of the Stoke’s coefficient for the cylinder in simulation units can be calculated to be Γsim= 55 s.u. Since our mass-spring system consists of 500 beads, each bead can be assigned a viscous Stoke’s coefficient of Γsim/(no. of beads) = 55/500 s.u. = 0.11 s.u.

#### e. Time

We can construct a unit of time using the drag coefficient, Γ, neutral length of the *Hydra*, L_0_ and the elastic energy stored upon strain of ε = 1 as

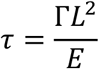

Having fixed the Stoke’s coefficient from the previous steps, we are now in a position to calculate a correspondence between the real time units and the simulation time units. Since all the parameters are known, we have

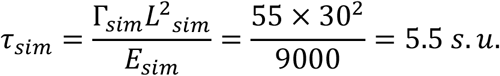

This must be equal to the real time units

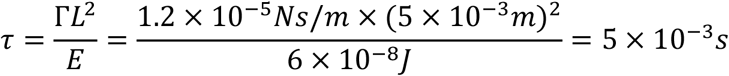

Comparing these two yields

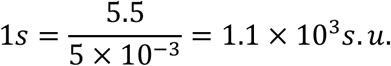

We have explicitly shown (see next section) that keeping the quantity Γ_sim_ Lsim^2^/E_sim_ constant yields the same relaxation behavior of the model.

The height of the basal disc from the floor versus time plots for ε = 0.2 for three different values of Γ_sim_ such that the value Γ_sim_ Lsim^2^/E_sim_ remains constant (with corresponding changes in E_sim_). We can see that the evolution in time is exactly the same, justifying its choice as the experimental time scale.

#### f. Estimation of E_Threshold_

We estimate the threshold energy required to overcome both viscous drag and gravitation by first performing a continuous integration of the drag force (per bead) along the path traced by each bead as the model-Hydra evolves in time from its configuration at the end of the contraction to its final configuration (inverted position). Since only the model-Hydras with α > 1 are able to reach this final configuration, we perform this integration for α = 3. The sum of all such integrals (over all 500 beads which constitute the cylinder) gives us the total drag energy dissipated from initial (after contraction) to final configuration. We must add to this the gravitational potential energy of the final configuration (∑m_i_gh_i_), which is obtained by summing over the gravitational potential energies of each bead in their final configuration. E_Threshold_ can then be formally written as:

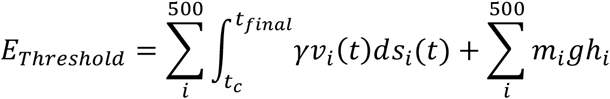

Here g’ is the effective the gravitational acceleration used for calculation. It is only a small fraction of the actual gravitational acceleration (g) to account for buoyancy resulting from density difference between *Hydra* tissue and water.

## Acknowledgments

We wish to thank Dr Sudhakaran Prabhakaran and Dr Sanjay Sane for useful comments on the manuscript. We thank Yashodeep Matange and Kanchan Sharma for help with animations and Dr Arpita Roychoudhury, Jitesh Seth and Adarsh Kumar for assistance with experiments. This work was supported by a grant from the Centre of Excellence in Epigenetics program (BT/01/COE/09/07) of the Department of Biotechnology, Government of India and the JC Bose National Fellowship (SG). We have used the computer cluster obtained using a grant from the Department of Biotechnology (BT/PR16542/BID/7/654/2016) to AC. AC acknowledges funding by DST Nanomission, India, the Thematic Unit Program (Grant No. SR/NM/TP-13/2016) and DST grant MTR/2019/000078. Authors acknowledge funding from IISER Pune - intramural (DS, SR, SG, SP and AC); Department of Biotechnology postdoctoral fellowship (PCR), the Wellcome Trust-Department of Biotechnology India Alliance for Intermediate Fellowship (500172/Z/09/Z) (SP) and Early Career Fellowship (IA/E/16/1/503057) (PCR); Department of Science and Technology, Govt. of India SERB grant No EMR/2015/000018 (AC); fellowships from the University Grants Commission (UGC) (MU), EMBO Short-term fellowship and Infosys foundation for international travel support (MU), Kishore Vaigyanik Protsahan Yojana (KVPY) (SN). The electron microscopy imaging was performed at the Irving and Cherna Moskowitz Center for Nano and Bio-Nano Imaging at the Weizmann Institute of Science, Rehovot, Israel.

## Footnotes

To whom correspondence should be addressed.

## Source Data legends

**Source data 1:** This source data file contains all the Young’s modulus data generated by AFM measurement presented in Figure 2 and Figure S1 C & D.

**Source data 2:** This dataset contains the raw data and the fitted curve data for the change in bending energy in the head region in Figure 3 D. The fit parameters and statistics for the fit are also provided.

**Source data 3:** This source data file contains all the datasets relating to model-*hydra* simulations and *hydra* density calculations presented in Figure 5 and Figure S2.

**Source data 4:** This source data file contains all the datasets pertaining to quantification of somersaults performed by *Hydra* upon partial amputation and Dipyridyl treatment from Figure 6 and Figure S6. It also contains AFM datasets upon Dipyridyl treated and nicked *Hydra*.

## References

Alexander, R. M. 2003. Principles of animal locomotion, Princeton University Press.

Akhtar, R., Sherratt, M. J., Cruickshank, J. K. & Derby, B. 2011. Characterizing the elastic properties of tissues. Materials Today, 14, 96–105.

Anderson, E. J. & Demont, M. E. 2000. The mechanics of locomotion in the squid Loligo pealei: locomotory function and unsteady hydrodynamics of the jet and intramantle pressure. Journal of Experimental Biology, 203, 2851–2863.

Aufschnaiter, R., Wedlich-SÖldner, R., Zhang, X. & Hobmayer, B. 2017. Apical and basal epitheliomuscular F-actin dynamics during Hydra bud evagination. Biology Open, 6, 1137–1148.

Aufschnaiter, R., Zamir, E. A., Little, C. D., Özbek, S., Munder, S., David, C. N., Li, L., Sarras, M. P. & Zhang, X. 2011. In vivo imaging of basement membrane movement: ECM patterning shapes Hydra polyps. Journal of Cell Science, 124, 4027–4038.

Barnes, H.A., Hutton, J.F. and Walters, K. 1989. An introduction to rheology (Vol. 3). pp. 41.Elsevier.

Biewener, A. A. 1990. Biomechanics of mammalian terrestrial locomotion. Science, 250, 1097–1103.

Bode, H. R. 1996. The interstitial cell lineage of hydra: a stem cell system that arose early in evolution. Journal of Cell Science, 109, 1155–1164.

Bode, H. R., Gee, L. W. & Chow, M. A. 1990. Neuron differentiation in hydra involves dividing intermediates. Developmental Biology, 139, 231–243.

Bond, C. & Harris, A. K. 1988. Locomotion of sponges and its physical mechanism. Journal of Experimental Zoology, 246, 271–284.

Bray, D. 2000. Cell movements: from molecules to motility, pp. 8 & 18. Garland Science.

Carter, J. A., Hyland, C., Steele, R. E. & Collins, E.-M. S. 2016. Dynamics of mouth opening in Hydra. Biophysical Journal, 110, 1191–1201.

Cavagna, G. A., Heglund, N. C. & Taylor, C. R. 1977. Mechanical work in terrestrial locomotion: two basic mechanisms for minimizing energy expenditure. American Journal of Physiology-Regulatory, Integrative and Comparative Physiology, 233, R243–R261.

Davis, L. E., Burnett, A. L., Haynes, J. F., Osborne, D. G. & Spear, M. L. 1968. Histological and ultrastructural study of the muscular and nervous systems in Hydra. II. Nervous system. Journal of Experimental Zoology, 167, 295–331.

Deutzmann, R., Fowler, S., Zhang, X., Boone, K., Dexter, S., Boot-Handford, R., Rachel, R. & Sarras, M. 2000. Molecular, biochemical and functional analysis of a novel and developmentally important fibrillar collagen (Hcol-I) in hydra. Development, 127, 4669–4680.

Dhont, J. K. 1996. An introduction to dynamics of colloids, Elsevier.

Dickinson, M. H., Farley, C. T., Full, R. J., Koehl, M., Kram, R. & Lehman, S. 2000. How animals move: an integrative view. science, 288, 100–106.

Dupre, C. & Yuste, R. 2017. Non-overlapping neural networks in Hydra vulgaris. Current Biology, 27, 1085–1097.

Engler, A. J., Sen, S., Sweeney, H. L. & Discher, D. E. 2006. Matrix elasticity directs stem cell lineage specification. Cell, 126, 677–689.

Ewer, R. & Fox, H. M. 1947. On the Functions and Mode of Action of the Nematocysts of Hydra. Proceedings of the Zoological Society of London. Wiley Online Library, 365–376.

Galliot, B. 2000. Conserved and divergent genes in apex and axis development of cnidarians. Current Opinion in Genetics & Development, 10, 629–637.

Gemmell, B. J., Colin, S. P., Costello, J. H. & Dabiri, J. O. 2015. Suction-based propulsion as a basis for efficient animal swimming. Nature Communications, 6, 1–8.

Gray, J. 1933. Studies in animal locomotion: I. The movement of fish with special reference to the eel. Journal of experimental biology, 10, 88–104.

Han, S., Taralova, E., Dupre, C. & Yuste, R. 2018. Comprehensive machine learning analysis of Hydra behavior reveals a stable basal behavioral repertoire. Elife, 7, e32605.

Hansen, P., Hassenkam, T., Svensson, R. B., Aagaard, P., Trappe, T., Haraldsson, B. T., Kjaer, M. & Magnusson, P. 2009. Glutaraldehyde cross-linking of tendon—mechanical effects at the level of the tendon fascicle and fibril. Connective Tissue Research, 50, 211–222.

Haynes, J. F., Burnett, A. L. & Davis, L. E. 1968. Histological and ultrastructural study of the muscular and nervous systems in Hydra. I. The muscular system and the mesoglea. Journal of Experimental Zoology, 167, 283–293.

Holstein, T. W., Hobmayer, E. & David, C. N. 1991. Pattern of epithelial cell cycling in hydra. Developmental biology, 602–611.

Hu, W., Lum, G. Z., Mastrangeli, M. & Sitti, M. 2018. Small-scale soft-bodied robot with multimodal locomotion. Nature, 554, 81–85.

Lenhoff, H. 2013. Hydra: research methods, Springer Science & Business Media.

Markowitz, J. & Herr, H. 2016. Human leg model predicts muscle forces, states, and energetics during walking. PLoS computational biology, 12, e1004912.

Martínez, D., Iñiguez, A., Percell, K., Willner, J., Signorovitch, J. & Campbell, R. 2010. Phylogeny and biogeography of Hydra (Cnidaria: Hydridae) using mitochondrial and nuclear DNA sequences. Molecular Phylogenetics and Evolution, 57, 403–410.

Matsumoto, G. 1991. Swimming movements of ctenophores, and the mechanics of propulsion by ctene rows. Hydrobiologia. Springer, 319–325.

Park, J. S., Chu, J. S., Tsou, A. D., Diop, R., Tang, Z., Wang, A. & Li, S. 2011. The effect of matrix stiffness on the differentiation of mesenchymal stem cells in response to TGF-β. Biomaterials, 32, 3921–3930.

Reddy, P. C., Barve, A. & Ghaskadbi, S. 2011. Description and phylogenetic characterization of common hydra from India. Current Science, 101, 736–738.

Reddy, P. C., Unni, M. K., Gungi, A., Agarwal, P. & Galande, S. 2015. Evolution of Hox-like genes in Cnidaria: Study of Hydra Hox repertoire reveals tailor-made Hox-code for Cnidarians. Mechanisms of Development, 138, 87–96.

Roberts, T. J. 2016. Contribution of elastic tissues to the mechanics and energetics of muscle function during movement. Journal of Experimental Biology, 219, 266–275.

Sarras JR, M. P. 2012. Components, structure, biogenesis and function of the Hydra extracellular matrix in regeneration, pattern formation and cell differentiation. International Journal of Developmental Biology, 56, 567–576.

Sarras JR, M. P., Meador, D. & Zhang, X. 1991. Extracellular matrix (Mesoglea) of Hydra vulgaris: II. Influence of collagen and proteoglycan components on head regeneration. Developmental Biology, 148, 495–500.

Shimizu, H., Aufschnaiter, R., Li, L., Sarras JR, M. P., Borza, D.-B., Abrahamson, D. R., Sado, Y. & Zhang, X. 2008. The extracellular matrix of hydra is a porous sheet and contains type IV collagen. Zoology, 111, 410–418.

Shimizu, H., Zhang, X., Zhang, J., Leontovichi, A., Fei, K., Yan, L. & Sarras, M. P. 2002. Epithelial morphogenesis in hydra requires de novo expression of extracellular matrix components and matrix metalloproteinases. Development, 129, 1521–1532.

Shostak, S., Patel, N. & Burnett, A. 1965. The role of mesoglea in mass cell movement in Hydra. Developmental biology, 12, 434–450.

Siegel, R. C., Pinnell, S. R. & Martin, G. R. 1970. Cross-linking of collagen and elastin. Properties of lysyl oxidase. Biochemistry, 9, 4486–4492.

Tao, N., Lindsay, S. & Lees, S. 1992. Measuring the microelastic properties of biological material. BiophysicalJjournal, 63, 165.

Taylor, C. R. & Heglund, N. C. 1982. Energetics and mechanics of terrestrial locomotion. Annual Review of Physiology, 44, 97–107.

Trembley, A. 1744. Mémoires pour servir à l’histoire d’un genre de polypes d’eau douce, à bras en forme de cornes, Chez Jean & Herman Verbeek.

Wood, R. L. 1979. The fine structure of the hypostome and mouth of hydra. Cell and Tissue Research, 199, 319–338.

Zhang, X., Boot-Handford, R. P., Huxley-Jones, J., Forse, L. N., Mould, A. P., Robertson, D. L., Athiyal, M. & Sarras, M. P. 2007. The collagens of hydra provide insight into the evolution of metazoan extracellular matrices. Journal of Biological Chemistry, 282, 6792–6802.

